# STK25 inhibits PKA signaling by phosphorylating PRKAR1A

**DOI:** 10.1101/2022.01.27.478084

**Authors:** Xiaokan Zhang, Bryan Z. Wang, Michael Kim, Trevor R. Nash, Bohao Liu, Jenny Rao, Roberta Lock, Manuel Tamargo, Rajesh Kumar Soni, John Belov, Eric Li, Gordana Vunjak-Novakovic, Barry Fine

**Author notes:** **Corresponding Author**, Barry Fine MD PhD, Department of Medicine, Division of Cardiology, 622 West 168^th^ Street, PH8-405B, New York, NY 10032.

## Abstract

In the heart, Protein Kinase A (PKA) is critical for activating calcium handling and sarcomeric proteins in response to beta adrenergic stimulation leading to increased myocardial contractility and performance. The catalytic activity of PKA is tightly regulated by regulatory subunits which inhibit the catalytic subunit until released by cAMP binding. Phosphorylation of Type II regulatory subunits promotes PKA activation, however the role of phosphorylation in Type I regulatory subunits remain uncertain. Here we utilized human induced pluripotent stem cell cardiomyocytes (iPSC-CM) to identify STK25 as a kinase of the Type Ia regulatory subunit PRKAR1A. Phosphorylation of PRKAR1A led to inhibition of PKA kinase activity and increased binding to the catalytic subunit in the presence of cAMP. *Stk25* knockout in mice diminished Prkar1a phosphorylation, increased Pka activity and augmented contractile response to beta adrenergic stimulation. Together, these data support STK25 as a negative regulator of PKA signaling through phosphorylation of PRKAR1A.

## Introduction

Second messengers are key relays in the transduction of external cues into intracellular signaling cascades. cAMP is a prototypical second messenger generated downstream from G protein coupled receptors and is involved in integrating a diverse set of signaling pathways (Gancedo, 2013). The classic and most widely studied effector of cAMP is PKA and it has been broadly implicated in a diverse set of cellular processes including the cell cycle, proliferation, cytoskeletal dynamics, ion flux and beta adrenergic signaling (Tasken and Aandahl, 2004; Taylor et al., 2012). Specificity of signaling in a system with such a broad set of substrates relies on careful regulation of both cAMP metabolism as well as compartmentalization of PKA through a series of A kinase anchoring proteins (AKAP) (Colledge and Scott, 1999). These AKAPs bind to the regulatory subunits of PKA and direct its activity to a discrete set of effectors and substrates, thereby allowing for enhanced spatiotemporal control over PKA and the pathways it activates.

The PKA holoenzyme is comprised of a regulatory subunit dimer and two catalytic units (Kim et al., 2007). The main function of the regulatory subunit is to inactivate kinase activity in the absence of cAMP. There are two classes of regulatory subunits, Type I and Type II, each with an alpha and beta isoform. Binding of cAMP to the C terminal tandem cAMP binding domains of the regulatory subunits releases the catalytic domain, allowing for kinase activity (Barradeau et al., 2002). Type II regulatory subunits have a PKA substrate motif in the N -terminus (RRXS), which is phosphorylated by the catalytic subunit upon binding to cAMP, enhancing its release and activation (Taylor et al., 1990). They also are the primary target of AKAPs, binding at nanomolar affinities (Colledge and Scott, 1999). The regulation of Type I subunits however remains less well understood. They contain a pseudo-substrate (RRXxA/G) sequence that cannot be phosphorylated by the catalytic subunit. Although phosphorylation of the Type I subunits has been reported as part of large phosphoproteomic projects, their role in regulation of PKA activity has not been demonstrated *in vivo* (Daub et al., 2008; Olsen et al., 2006; Olsen et al., 2010; Oppermann et al., 2009; Zhou et al., 2013). Furthermore, the vast majority AKAP proteins do not interact with Type I subunits, and the dual specificity AKAP proteins that do interact with Type I subunits do so with significantly lower affinities compared to Type II subunits.

In the heart, PKA mediates beta adrenergic activation of the physiologic flight or fight response, modulating both heart rate and myocardial contractility (Wang et al., 2018). Substrates of PKA in the cardiomyocyte include troponin (TNNT2), myosin binding protein c (MYBPC3), phospholamban (PLN), and ryanodine receptor (RYR2) (Marks, 2013). Acute activation of PKA in hyperadrenergic states leads to improved calcium flux and myocardial performance and is necessary for increases in stroke volume to meet cardiac output demands. In heart failure, chronic beta-adrenergic stimulation from high levels of catecholamines results in compensatory downregulation of beta adrenergic signaling to leading diminished phosphorylation of several PKA substrates (Bristow et al., 1982; Piacentino et al., 2003). Long term activation of Pka was shown to be deleterious in mouse models and is thought to underly the maladaptive remodeling of chronic beta-adrenergic activation in heart failure (Antos et al., 2001).

STK25 is a stress response kinase in the sterile-20 (Ste20) kinase superfamily and has been shown through a series of *in vivo* models to impact glucose utilization, insulin homeostasis, and lipid metabolism (Amrutkar et al., 2015a; Amrutkar et al., 2015b; Amrutkar et al., 2016a; Amrutkar et al., 2016b; Cansby et al., 2013; Chursa et al., 2017; Nerstedt et al., 2012; Nunez-Duran et al., 2017; Sutt et al., 2018). Overexpression of STK25 leads to steatosis and was shown to aggravate atherosclerosis in a PCSK9 gain of function mouse model (Cansby et al., 2018). Recently, STK25 was shown to interact with the Hippo signaling via phosphorylation of LATS1/2 and/or regulation of SAV1-STRIPAK (Bae et al., 2020; Lim et al., 2019). In this study, we use a combination of *in vitro* and *in vivo* cardiomyocytes models to demonstrate that STK25 phosphorylates the Type Ia regulatory subunit of PKA, PRKAR1A. This leads to increased inhibition of PKA signaling, revealing a new regulatory mechanism that can control PKA activity and attenuate cAMP mediated increases in cardiac contractile function.

## Methods

### Patient samples

Patients with advanced HF were recruited at Columbia University Medical Center and heart tissue at the time of heart transplant was collected from the left ventricle. Control myocardial samples were obtained from the National Disease Research Interchange (https://ndriresource.org/ats) and were comprised of de-identified specimens collected from non-failing hearts determined to be unusable for cardiac transplantation due to non-cardiac donor issues but without evidence or knowledge of underlying cardiac disease. The study was approved by the Institutional Review Board of Columbia University (IRB# AAAR0055). All patients provided written informed consent before inclusion into the study.

### Cell culture

HEK293T cells (ATCC Cat# CRL-3216, RRID:CVCL_0063) were cultured in high glucose (4.5g/L) DMEM supplemented with 10% FBS, penicillin and streptomycin, and grown in a CO_2_ incubator maintained at atmospheric oxygen levels and 5% CO_2_. Human induced pluripotent stem cells (hiPSC) were obtained through Material Transfer Agreements from Bruce Conklin, Gladstone Institute (WTC cell line). hiPSCs were expanded on growth factor reduced Matrigel-coated plates (Corning) in mTeSR plus medium (Stemcell technologies) containing mTeSR plus supplement (Stemcell technologies), 50U penicillin and 50U streptomycin. The cell culture medium was changed every other day, and cell passaged upon reaching 70% confluence. During the first 24 hrs after passaging, 5 μM Y-27632 dihydrochloride (Tocris, 1254) was supplemented to culture medium.

### CRISPR-Cas9

CRISPR-Cas9 Genome editing was used to generate a homozygous knockout of *STK25* in WTC iPSCs following the manufacturer’s protocol (ORIGENE) with gRNA vector (KN203215G). Single-cell clones were expanded and confirmation of homozygous editing was determined by sanger sequencing of gDNA and protein analysis by western blot.

### iPS Cardiomyocyte Differentiation

Cardiac differentiation of iPSC’s was initiated in confluent monolayers two days after replating at a density of 200,000/cm^2^ (24hrs in mTeSR plus medium with 5μM Y-27632 dihydrochloride, followed by 24hrs in mTeSR plus medium without Y-27632). On day 0 of differentiation, media was changed to cardiac differentiation medium (CDM, consisting of RPMI-1640, Life Technologies), 500μg/ml human recombinant albumin (Sigma), 213μg/ml L-ascorbic acid (Sigma), 50U penicillin and 50U streptomycin) with 3-6 μM CHIR (4423, Tocris). 2 days after initial CHIR addition, media was changed to CDM with 2 μM Wnt-C59 (5148, Tocris). From day 4 onwards, media was changed fresh CDM every other day until cells start contracting. Media was then switched to RPMI-B27 (consisting of RPMI-1640, 1x B27 supplement (Life Technologies), 213μg/mL L-ascorbic acid, 50U penicillin and 50U streptomycin). Cardiomyocytes were characterized by flow cytometry using the cardiomyocyte-specific marker TNNT2 (BD Biosciences Cat# 565744, RRID:AB_2739341). Differentiation typically resulted in cell populations containing 80–90% TNNT2-positive cells at day 12.

### PKA activity assay

PKA activity assay was performed using whole cell lysate from iPSC-CMs treated with either PBS or 10μM forskolin (Sigma-Aldrich, F6886) for 30min at 37°C prior to assay, or immunoprecipitated type I and II PKA holoenzymes from iPSC-CMs whole cell lysate or whole heart lysates from *Stk25^+/+^* or *Stk25^-/-^* mice following the manufacturer’s protocol (EIAPKA, Invitrogen). Briefly, PKA standards or diluted samples and reconstituted ATP were added into wells of PKA substrate plate. Phospho-PKA substrate antibody and Goat anti-Rabbit IgG HRP conjugate antibody were added into the wells per manufacturer protocol. After incubation TMB substrate was added into each well, and the plate was incubated for 30 min at room temperature. Stop solution was added, and absorbance at 450nm was analyzed in a 96-plate reader.

### Global Phosphoproteomics analysis

Sorted *STK25^+/+^* and *STK25^-/-^* iPSC-CMs were lysed/homogenized by bead-beating in 8 M urea, 1%SDS and 200 mM EPPS (pH 8.5), protease, and phosphatase inhibitors. Lysates were cleared by centrifugation at 21,000 g for 30 min at 4 °C, and protein concentration was measured by BCA. Proteins were reduced with 5 mM TCEP, alkylated with 10 mM iodoacetamide (IAA), and quenched with 10 mM DTT. A total of 500 μg of protein was chloroform-methanol precipitated. Protein was reconstituted in 200 mM EPPS (pH 8.5) and digested by Lys-C overnight and trypsin for 6 h, both at a 1:50 protease-to-peptide ratio. Digested peptides were quantified using a Nanodrop at 280 nm, and 200 μg of peptide from each sample were labeled with 800 μg TMT reagent using a 10-plex TMT kit (Navarrete-Perea et al., 2018). TMT labels were checked, 100 ng of each sample was pooled and desalted and analyzed by short SPS-MS3 method, and using normalization factor samples were bulk mixed at 1:1 across all channels and desalted using a 200 mg Sep-Pak solid-phase extraction column and dried using vacuum centrifugation.

Desalted isobaric labeled peptides were enriched for phospho-peptides using a mixture of MagReSyn Ti-IMAC and Zr-IMAC resins according ReSyn Bioscience instructions. In brief, 1.4 mg of labeled peptide were resuspended in 1 ml of binding buffer (80% Acetonitrile, 1M glycolic acid and 5% TFA) and incubated with equilibrated 150 μl (75 μl of each Ti-IMAC and Zr-IMAC) resins at room temperature for 30 min, and the resin was washed 3 three times to remove the unbound, non-phosphorylated peptides. Phospho-peptides were eluted using 1% ammonium hydroxide. The enriched phospho-peptides were further fractionated in eight fractions using Pierce™ High pH Reversed-Phase Peptide Fractionation Kit and each fraction dried down in a speed-vac.

The isobaric labeled dried phospho-peptides were resuspended in 10 μl of (3% acetonitrile/ 0.1% formic acid), and analyzed on an Orbitrap Fusion mass spectrometer coupled to a Dionex Ultimate 3000 (ThermoFisher Scientific) using the MSA-SPS-MS3 and NL SPS-MS3 method (Jiang et al., 2017). Peptides were separated on an EASY-Spray C18 50cm column (Thermo Scientific). Peptide elution and separation were achieved at a non-linear flow rate of 250 nl/min using a gradient of 5-30% of buffer B (0.1% (v/v) formic acid, 100% acetonitrile) for 110 minutes with a temperature of the column maintained at 50 °C during the entire experiment. For both methods, MS1 data were collected using the Orbitrap (120,000 resolution; maximum injection time 50 ms; AGC 4×10^5^). Determined charge states between 2 and 5 were required for sequencing and a 45 s dynamic exclusion window was used. Data-dependent top10 MS2 scans were performed in the ion trap with collision-induced dissociation (CID) fragmentation (Turbo; NCE 35%; maximum injection time 60ms; AGC 5×10^4^). MS3 quantification scans were performed using the multi-notch MS3-based TMT method (ten SPS ions; 50,000 resolution; NCE 65% for MSA-SPS-TMT and 38% for NL-SPS-TMT maximum injection time 105 ms; AGC 1×10^5^) using the Orbitrap.

Raw mass spectrometric data were analyzed using Proteome Discoverer 2.2 to perform database search and TMT reporter ions quantification. TMT tags on lysine residues and peptide N termini (+229.163 Da) and the carbamidomethylation of cysteine residues (+57.021 Da) was set as static modifications, while the oxidation of methionine residues (+15.995 Da), deamidation (+0.984) on asparagine and glutamine and phosphorylation (+79.966) on serine, threonine, and tyrosine were set as a variable modification. Data were searched against a UniProt Human database with peptide-spectrum match (PSMs) and protein-level FDR at 1% FDR. The signal-to-noise (S/N) measurements of each protein normalized so that the sum of the signal for all proteins in each channel was equivalent to account for equal protein loading. Phospho-peptides identification and quantification were imported into Perseus (Tyanova et al., 2016) for t-test statistical analysis (FDR<0.05) to identify phospho-peptides demonstrating statistically significant changes in abundance. Pathway analysis were performed using Ingenuity Pathway Analysis (Qiagen).

### Animal studies

*Stk25* knockout mice were generated by CRISPR/Cas9 mediated genome engineering in C57BL/6J background (Jackson Laboratory, Bar Harbor, Maine). Exon 3-5 of Stk25 gene were deleted with two CRISPR guides from Synthego (sgRNA-stk25-7367 and sgRNA-stk25-10150). Both guides were mixed with IDT Cas9V3 protein to form RNP, which was further injected into the pronuclei of fertilized C57BL/6J eggs to generate knockout founders. Genotyping was performed to show the *Stk25* knockout allele. Further breeding generated homozygous *Stk25^-/-^* mice and protein immunoblotting was performed to confirm an absence of Stk25. The protocol for all mouse experiments (AABC1503) was approved by the Columbia University Institutional Animal Care and Use Committee.

Cardiac function of 20 week and 52 week old mice was assessed by echocardiography by the Columbia University Mouse Imaging Core Facility (imaged by Visulasonics VEVO 3100 High Frequency Ultrasound imaging system and analyzed by Vevo LAB software). For 20 week old mice, cardiac function was recorded at baseline and 3 minutes after administration of the β-adrenergic receptor agonist isoproterenol (0.2ug/g, i.p.). Systolic function parameters including ejection fraction (EF, %), fractional shortening (FS, %), left ventricular end systolic diameter (LVESD, mm) and left ventricular end diastolic diameter (LVEDD, mm) were measured in the two-dimensional parasternal short-axis imaging plane of M-mode tracings close to the papillary muscle level.

### iPS Cardiomyocyte sorting

Cardiomyocytes were pretreated with 10μM of Y-27632 in B27 culture medium for 6 hrs, then digested with cell dissociation buffer (containing Hank’s buffered saline solution, 100 Units/ml collagenase, 10μM Y-27632) at room temperature overnight. Cells were collected and centrifuged at 300g for 5 min, and then resuspended in cell sorting buffer (CSB, containing Hank’s buffered saline solution, 20mM HEPES, 5% FBS, 10 units/ml Turbo DNase). The cell pellet was resuspended with anti-CD172a/b (SIRPα/β, 423107, Biolegend) and anti-CD90 (Thy-1, 11-0909-42, Invitrogen) antibodies in CSB to stain for 20 min at 4°C in the dark. Cells were washed 3 times with CSB and DAPI staining was added. Cardiomyocytes were sorted from the SIRPA+ and CD90-population and then plated on matrigel coated plates in B27 medium with 5μM Y-27632. The following day, medium was replaced with fresh B27, and cardiomyocytes generally started to beat in 2-5 days.

### RNA Isolation, Sequencing, and Analysis

RNA was extracted using the RNeasy Mini kit (Qiagen #74004) along with on-column DNAse digestion (Qiagen #79254). cDNA libraries were generated using the Clontech Ultra Low v4 kit followed by NextaraXT DNA Library Prep Kit, then sequenced on an Illumina NovaSeq 6000. Paired-end 100-bp sequenced reads were analyzed as follows: RTA (Illumina) software was used for base calling and bcl2fastq2 (version 2.19) for converting BCL to fastq format, coupled with adaptor trimming. A pseudoalignment to a kallisto index created from transcriptomes (build GRCh38) using kallisto (0.44.0). After pseudoalignment the R package DESeq2 (1.28.1) was used to normalize the count matrix and calculate differentially expressed genes. Geneset enrichment analysis was performed using GSEA software (4.10.0) per documentation (Mootha et al., 2003; Subramanian et al., 2005).

### siRNA and vector transfection

Cells were transfected using lipofectamine 3000 (Invitrogen) following the manufacturer’s protocol with siRNA targeting STK25 (On-TargetPlus siRNA, Horizon Discovery, L-004873-00-0050) or siRNA targeting PRKAR1A (EHU071341, Sigma) for protein knockdown or non-targeting control siRNA (ON-TARGETplus non-targeting pool, Horizon Discovery D-001810-10-50) as scramble control.

For heterologous expression studies, cells were transfected using lipofectamine 3000 (Invitrogen, L3000015) following the manufacturer’s protocol. Vectors used include Flag-WT-STK25 vector (EX-M0142-M46, GeneCopoeia), Flag-K49R/T147A-STK25 vector (Vector Builder), Flag- empty vector (EX-NEG-M46, GeneCopoeia), V5-PRKAR1A (Vector Builder, VB200124-1141aes), V5-S77A/S83A-PRKAR1A (Vector Builder, VB200124-1126jtp), and V5-S77E/S83E-PRKAR1A (Vector Builder, VB200124-1127pst).

### qRT-PCR assays

Equivalent amounts (2μg) of purified RNA were used as a template to synthesize cDNA using oligo-d(T) primers and SuperScript™ III First-Strand Synthesis SuperMix (Invitrogen, 18080400). *STK25* was quantified by real-time PCR using Fast SYBR Green mixture (Life Technologies, 4385612) and was carried out on Applied Biosystems Step One Plus. Relative levels were calculated using ΔΔCτ method. Data analysis was carried out using the fold change normalized to GAPDH gene expression.

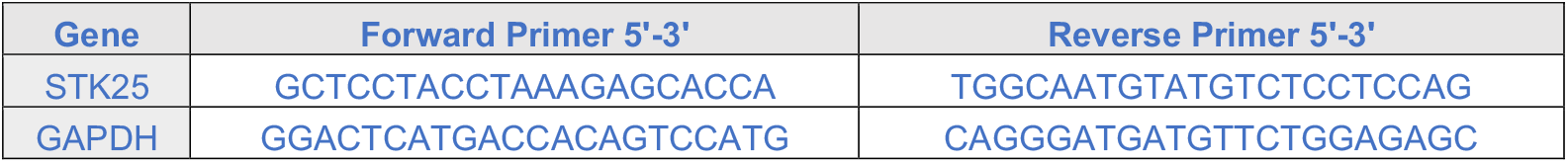

### Immunoprecipitation and immunoblot analysis

300 μg of total protein from whole cell lysate were used in immunoprecipitation experiments. The extract was incubated with anti-Flag affinity gel (Sigma-Aldrich Cat# A2220, RRID:AB_10063035), anti-V5 affinity gel (Sigma-Aldrich Cat# A7345, RRID:AB_10062721), anti-PRKAR1A (Abcam Cat# ab139695, RRID:AB_2893184) or anti-PRKAR2A (ProteinTech Cat# 10142-2-AP) overnight at 4°C. The affinity gel beads were centrifuged at 8000xg for 1min, washed three times in TBS buffer. The affinity gel beads were added with 4x Laemmli SDS sample buffer (NuPAGE), denatured at for 5min at 95°C and analyzed by SDS-PAGE and immunoblotting. For Western blotting antibodies include: HRP conjugated anti-GAPDH (Cell Signaling Technology Cat# 3683, RRID:AB_1642205), anti-STK25 (Abcam Cat# ab157188, RRID:AB_2725788), anti-Flag (Sigma-Aldrich Cat# F3165, RRID:AB_259529), anti-GM130 (Cell Signaling Technology Cat# 12480, RRID:AB_2797933), anti-PRKAR1A (Abcam Cat# ab139695, RRID:AB_2893184), anti-pS77 PRKAR1A (Abcam Cat#ab139682, RRID:AB_2904566), anti-pS83 PRKAR1A (Abcam Cat#ab154851, RRID:AB_2904567), anti-PRKAR2A (ProteinTech Cat# 10142-2-AP), anti-phospholamban (Cell Signaling Technology Cat# 14562, RRID:AB_2798511), anti-phospho-phospholamban -pS16/T17 (Cell Signaling Technology Cat# 8496, RRID:AB_10949102), anti-Ryanodine receptor 2 (Abcam Cat#ab196355, RRID:AB_2904568), anti-p-Ryanodine receptor 2-pS2808 (Abcam Cat# ab59225, RRID:AB_946327), anti-TnI (Cell Signaling Cat# 4002), anti-TnI-pS23/S24 (Cell Signaling Cat# 4004), anti-V5 (Sigma-Aldrich Cat# V8137, RRID:AB_261889), anti-PKA Catalytic subunit (Abcam Cat# ab26322, RRID:AB_2170049), HRP-conjugated anti-mouse (Cell Signaling Technology Cat# 7076, RRID:AB_330924) and HRP-conjugated anti-rabbit (Cell Signaling Technology Cat# 7074, RRID:AB_2099233) were used for detection.

### In vitro kinase assay

Recombinant protein STK25 (0.5μM, TP303215, OriGene) and PRKAR1A (1μM, TP303828 Origene) were mixed with *<u>in vitro</u>* kinase buffer (25mM Tris-HCl, 10mM beta-glycerophosphate, 10mM MgCl_2_, 20mM NaF, 2mM DTT, 1mM sodium orthovanadate, 1x Protease Inhibitor cocktail (Roche)) containing ATP (500uM) and incubated at 37°C for 30 min. In vitro kinase assay was terminated by adding Laemmli SDS sample buffer. Phosphorylation levels of PRKAR1A at S77 and S83 were analyzed by SDS-PAGE and immunoblotting.

### Real Time Glo cell viability assay

Cells were seeded into 96-well plates at a density of 3000 cells per well 24h before measurement. RealTime-Glo (Promega; G9712) reagent was added to each well per manufacturer protocol and measurement of luminescence was performed at selected time points on a BioTek Synergy JHTX with Gen5 data analysis software.

## Statistical Analysis

Statistical analyses were performed using Prism 8 (Graphpad Software). Results are presented as mean ± standard deviation. For comparisons between two groups, a two tailed unpaired t-test was used unless otherwise specific. Welch’s correction was utilized for two groups of unequal sizes. For multiple group comparisons, either one way or two-way (depending on the number of variables) ANOVA followed by multiple comparison post-hoc testing was performed as indicated using Prism 8. Notation in the text is as follows *p<0.05, **p<0.01, ***p<0.001 and ****p<0.0001

## Results

### STK25 inhibits PKA signaling

As part of an kinome knockout discovery project, CRISPR-Cas9 was used to generate a homozygous knockout of *STK25* in the wild type iPSC line WTC11 (Bruce Conklin, Gladstone Institute) (Hayashi et al., 2016), and cardiomyocytes were differentiated and characterized using an established protocol (Burridge et al., 2014). *STK25* expression in iPSC derived cardiomyocytes was found to be similar to *Stk25* expression to primary cardiomyocytes from adult mice (**Figure S1A and S1B**). To identify potential substrates of STK25 in cardiomyocytes, mass spectrometry based phosphoproteomics was performed with *STK25^+/+^* and *STK25^-/-^* cardiomyocytes (**Figure 1A**). Signaling pathways impacted by *STK25* were identified using Ingenuity Pathway Analysis (Qiagen). A decrease in Hippo signaling was observed in response to loss of STK25 confirming a prior study demonstrating STK25 activates the Hippo pathway (**Figure 1B**) (Lim et al., 2019). The most significant signaling change in response to loss of *STK25* was an upregulation of the PKA pathway. Several downstream substrates of PKA displayed increased phosphorylation including MYBPC3, TNNT2, RYR2, CACNA1C and GSK3α (**Table 1**). *STK25^-/-^* cardiomyocytes exhibited increased PKA kinase activity when stimulated with the adenylate cyclase agonist forskolin (**Figure 1C**). In heterologous overexpression studies in HEK293T cells, STK25 decreased forskolin induced PKA activity. This inhibition was dependent on the kinase domain of STK25 as overexpression of the kinase-dead mutant *STK25^K49R/T174A^* (Preisinger et al., 2004) did not inhibit PKA activity in response to forskolin (**Figure 1D)**.

**Figure 1.**
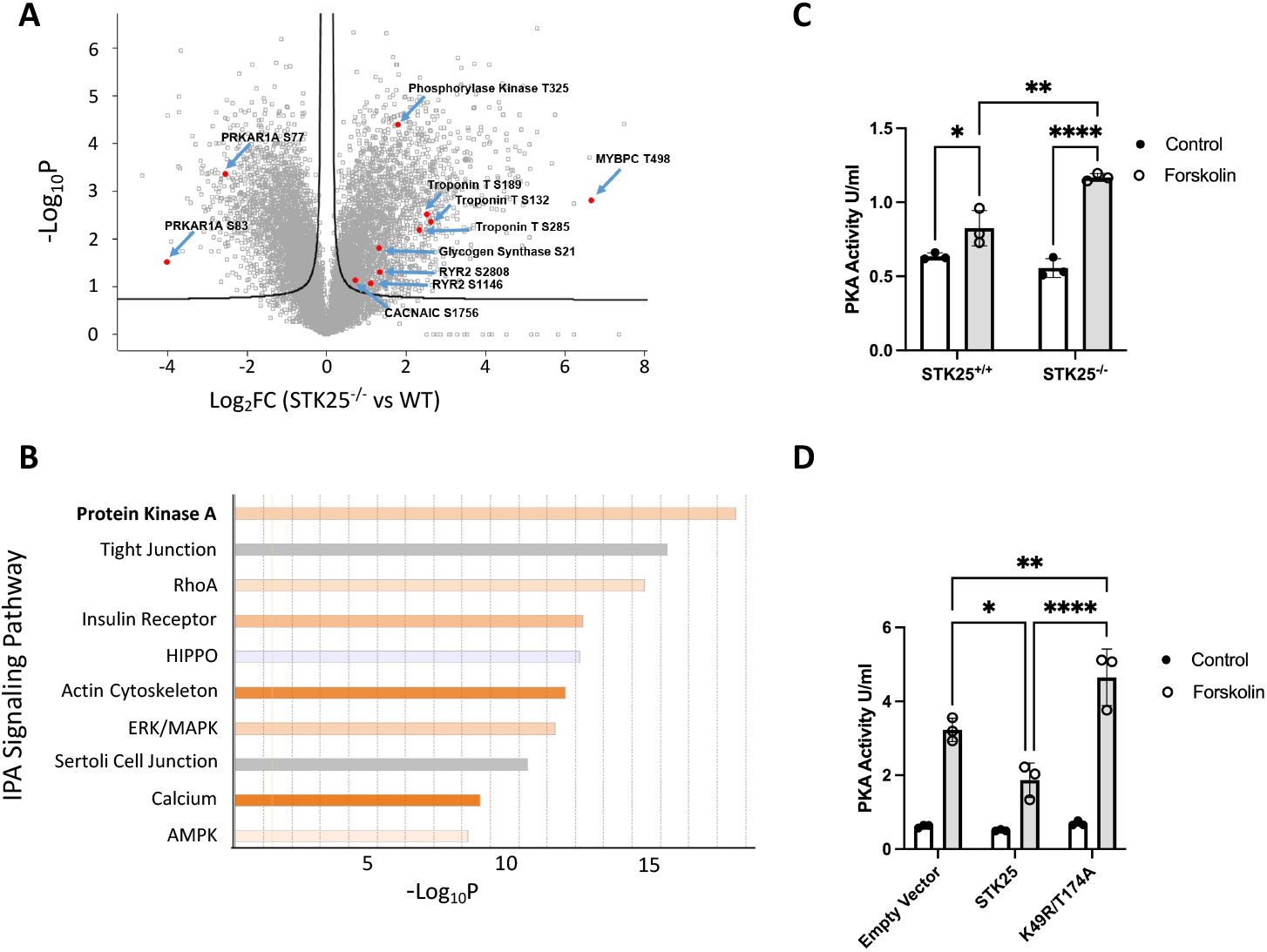
STK25 inhibits PKA activity. A) Differential phosphoproteomic spectra of *STK25^+/+^* and *STK25^-/-^* cardiomyocytes. Members of the PKA signaling pathway are highlighted. B) Ingenuity phosphoprotein pathway analysis. Orange indicates pathways upregulated in *STK25^-/-^* cardiomyocytes while blue indicates upregulation in *STK25^+/+^*. C) PKA activity in response to 10μM forskolin treatment for 30 minutes in *STK25^+/+^* and *STK25^-/-^* cardiomyocytes. n=3 for each condition. D) PKA activity in response to 10μM forskolin treatment for 30 minutes in HEK293T cells overexpressing either empty vector, wild type STK25 or kinase-dead K49R/T174A. n=3 for each condition. Bar graph data is represented as mean +/-SD, *p<0.05 **p<0.01, ****p<0.0001 using ANOVA and Tukey’s adjustment for multiple comparisons.

**Table 1:** PKA pathway phosphoproteomic changes in STK25^-/-^ compared to wild type. Fold change represents a ratio of STK25^-/-^/STK25^+/+^. Phosphorylation sites, description and accession numbers for each uniprot ID are listed. All changes met a significance threshold of an FDR corrected q value <0.05.

| Function |  | Protein Descriptions |  | Fold Change | Phosphorylation Sites | Protein Accessions |
| --- | --- | --- | --- | --- | --- | --- |
| PKA | Regulatory subunits | PRKAR1A | cAMP-dependent protein kinase type I-alpha regulatory subunit | 0.17 | S77, S83 | P10644 |
|  |  | PRKAR2A | cAMP-dependent protein kinase type II-alpha regulatory subunit | 3.03 | S78 | P13861 |
|  | Catalytic Subunit | PRKACA | cAMP-dependent protein kinase catalytic subunit alpha | 0.79 | S339 | P17612 |
| PKA targets | Contractility<br>Ca<sup>2+</sup> handling | MYBPC | Myosin-binding protein C, cardiac-type | 100.03 | T498 | Q14896 |
|  |  | Troponin | Troponin T, cardiac muscle | 6.18 | S132 | P45379 |
|  |  |  |  | 5.75 | S285 |  |
|  |  |  |  | 5.02 | S189 |  |
|  |  | RYR2 | Isoform 2 of Ryanodine receptor 2 | 2.51 | S2808 | Q92736 |
|  |  |  |  | 2.14 | S4368 |  |
|  |  | CACNA1C | Isoform 2 of Voltage-dependent L-type calcium channel subunit alpha-1C | 1.65 | S1756 | Q13936 |
|  | Energy Metabolism | Phosphorylase Kinase | Phosphorylase b kinase gamma catalytic chain | 3.48 | T325 | P15735 |
|  |  | Glycogen Synthase | Glycogen synthase kinase-3 alpha | 2.50 | S21 | P49840 |
|  | Gene Expression | CREB | Cyclic AMP-responsive element-binding protein 1 | 0.79 | S271 | P16220 |

### STK25 phosphorylates PRKAR1A

Two of the most downregulated phosphorylation sites in *STK25^-/-^* iPSC-CM were S77 and S83 of PRKAR1A raising the possibility that these sites were directly phosphorylated by STK25 (**Table 1**). PRKAR1A is the R1α member of the regulatory subunit isoforms of the PKA holoenzyme which binds to and inhibits the catalytic subunit of PKA (PRKACA) and has been shown to disengage in response to cAMP, leading to PKA activation (Bossis and Stratakis, 2004). We first confirmed the phosphoproteomic data by demonstrating decreased phosphorylation of S77 and S83 in *STK25^-/-^* cardiomyocytes by immunoblot protein analysis (**Figure 2A and S2A**). In order to determine if this observation was dependent on the kinase activity of STK25, STK25^K49R/T174A^ was expressed in *STK25^-/-^* iPSC-CMs and led to significantly decreased phosphorylation of S77 and S83 compared to overexpression of wild type STK25 (**Figure 2B and S2B**). Stimulation of PKA with forskolin led to diminished phosphorylation at both S77 and S83 in *STK25^+/+^* iPSC-CM but did not have significant effect on *STK25^-/-^* iPSC-CMs (**Figure 2C and S2C**).

**Figure 2.**
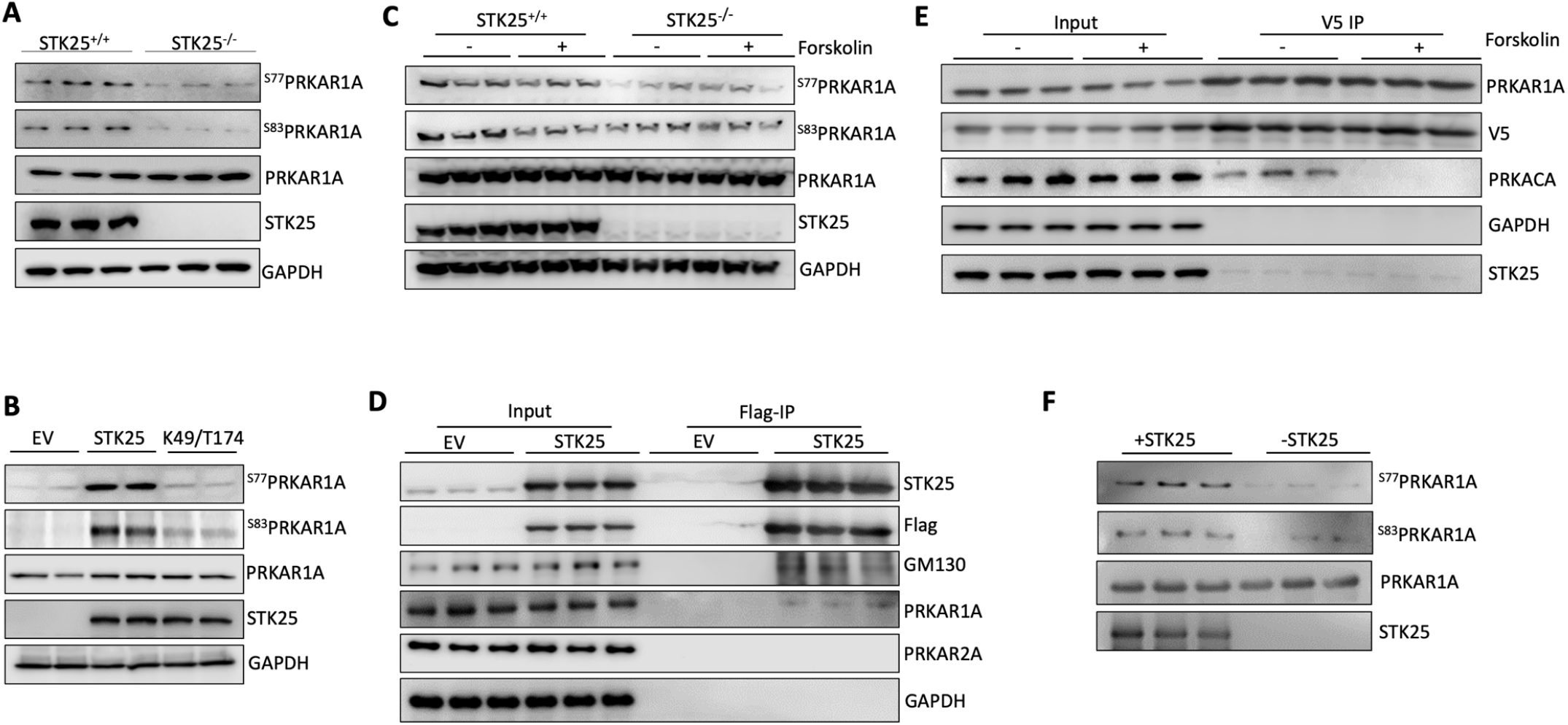
STK25 binds to and phosphorylates PRKAR1A. A) Immunoblot of PRKAR1A phospho-S77 and S83 in *STK25^+/+^* and *STK25^-/-^* cardiomyocyte protein lysates. B) Immunoblots of phosphor-S77 and S83 of PRKAR1A in *STK25^-/-^* cardiomyocytes transfected with empty vector (EV), wild type STK25 and kinase dead K49R/T174A STK25. C) Forskolin (10μM, 30 minutes) stimulated *STK25^+/+^* and *STK25^-/-^* cardiomyocytes immunoblotted for phosphorylation of PRKAR1A. D) Immunoprecipitation of Flag-STK25 expressed in HEK293T cells and immunoblotted for PRKAR1A, PRKA2A and GM130 (positive control binding partner). E) Co-immunoprecipitation of PRKAR1A-V5 with STK25 and PRKACA in HEK293T cells treated with forskolin (10μM, 30 minutes). F) *In vitro* kinase assay of purified STK25 and PRKAR1A, immunoblotted for phosphorylation of S77 and S83 of PRKAR1A.

Co-immunoprecipitations experiments demonstrated STK25 and PRKAR1A binding using epitope tagged protein in HEK293T cells (**Figure 2D**). Though forskolin inhibited binding between PRKAR1A and the catalytic subunit, PRKACA, it did not alter the association between STK25 and PRKAR1A (**Figure 2E**). Mutation of the kinase domain of STK25 also did not affect binding to PRKAR1A by immunoprecipitation (**Figure S2D**). Using purified protein in an *in vitro* kinase assay, STK25 was able to phosphorylate PRKAR1A at S77 and S83(**Figure 2F**).

### Phosphorylation of S77/S83 inhibits PKA activity

Phosphorylation at S77 and S83 on PRKAR1A have been described as part of large scale global phosphoproteomic experiments (Bian et al., 2014; Daub et al., 2008; Olsen et al., 2006; Oppermann et al., 2009) but their influence on PKA activity has not been characterized. In order to investigate the effect of pS77 and pS83 on the ability of PRKAR1A to inhibit PKA activity, phosphomimetic (S77E/S83E) and non-phosphomimetic (S77A/S83A) mutations were generated in PRKAR1A expression vectors. In HEK293T co-immunoprecipitation experiments, all three isoforms bind PRKACA similarly (**Figure S3A and S3B**). However, upon stimulation with forskolin the S77E/S83E mutant exhibited elevated binding to the catalytic subunit compared to the S77A/S83A and WT PRKAR1A, indicating that positive charges at this residue may interfere with release of PKA in response to cAMP (**Figure 3A and S3C)**. We then further tested this hypothesis by assessing PKA kinase activity *in vitro* in response to overexpression of these constructs in HEK293T cells. Though both WT and S77A/S83A PRKAR1A exhibited attenuated increases in induced PKA activity, expression of the S77E/S83E mutant exhibited no increase in PKA activity in response to forskolin (**Figure 3B**). Similarly, S77E/S83E PRKAR1A was able to inhibit metabolic activity of HEK293T in response to forskolin (**Figure 3C)** and in normal growth conditions (**Figure S3D)** to a significantly greater degree than either PRKAR1A or STK25 overexpression alone. In order to confirm that PRKAR1A is downstream of STK25, STK25 was overexpressed simultaneously with knock-down of PRKAR1A resulting in rescue of STK25 inhibition of PKA activity (**Figure 3D and Figure S3E**). Together these data demonstrate a new regulatory event where STK25 phosphorylation increases the ability of PRKAR1A to bind the catalytic subunit and inhibit PKA activity in response to cAMP (**Figure 3E)**.

**Figure 3.**
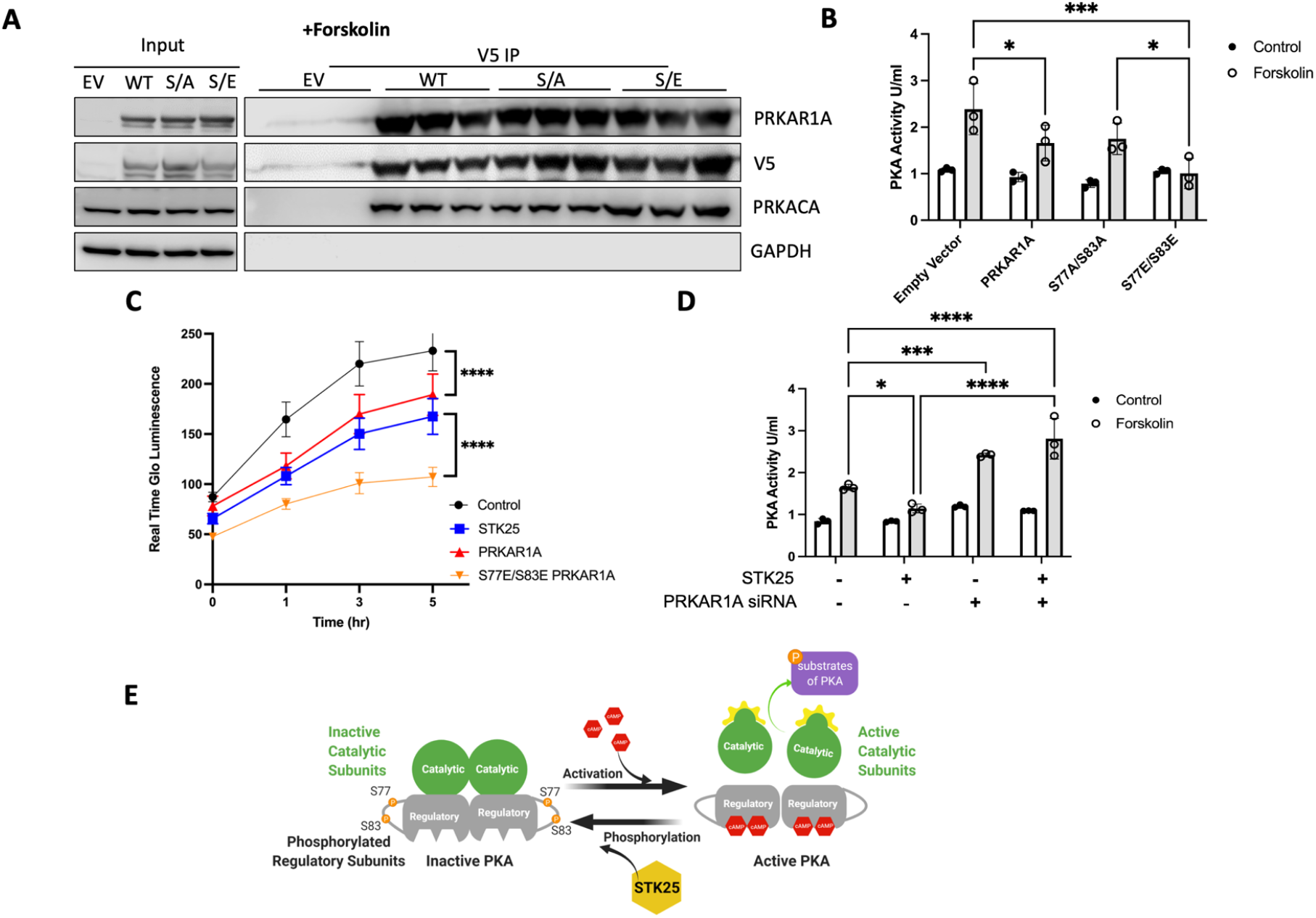
Phosphorylation of PRKAR1A inhibits PKA activity. A) Co-immunoprecipitations of V5 tagged PRKAR1A, S77A/S83A PRKAR1A or S77E/S83 PRKAR1A and immunoblotting for PRKARCA in HEK293T cells stimulated with forskolin (10μM, 30 minutes). B) PKA activity in HEK293T cells stimulated with forskolin (10μM, 30 minutes) and transfected with empty vector, wild type PRKAR1A, S77A/S83A PRKAR1A mutant or S77E/S83E PRKAR1A mutant as indicated. C) HEK293T cells transfected with the indicated vectors and assessed for growth by Real Time Glo for 5 hours after stimulation with forskolin (10μM). D) PKA activity in HEK293T cells stimulated with forskolin (10μM, 30 minutes) and either overexpressing STK25 and/or the siRNA of PRKAR1A. E) Model of STK25 downregulation of the PKA pathway through phosphorylation of PRKAR1A. For all graphs in this figure, n=3 for each condition, data presented as mean +/- SD *p<0.05, ***p<0.001, and ****p<0.0001 by ANOVA with Tukey’s adjustment for multiple comparisons.

Phosphorylation of PRKAR2A was also detected in our phosphoproteomic analysis however phosphorylation levels increased with the loss of STK25 (**Table 1)**. Despite this, given its role in PKA regulation in cardiomyocytes, we evaluated PKA holoenzyme activity from PRKAR1A and PRKAR2A immunoprecipitates. Loss of STK25 increased PKA activity from PRKAR1A complexes but not PRKAR2A complexes (**Figure S4A and S4B**). Furthermore, immunoprecipitation experiments did not demonstrate binding of PRKAR2A and STK25 (**Figure 2D**). Together these data do not support PRKAR2A as a downstream effector of STK25.

### *Stk25* loss increases PKA activity *in vivo*

In order to validate the STK25-PKA relationship *in vivo*, we generated an *Stk25^-/-^* mouse and investigated the impact of *Stk25* loss on the Pka pathway *in vivo*. Protein analysis demonstrated that loss of *Stk25* was accompanied by an increase in Prkar1a levels and a significant decrease in the ratio of pS77 and pS83 to total Prkar1a (**Figure 4A and S5A**). Whole heart lysates of *Stk25^-/-^* mice displayed increased levels of Pka kinase activity compared to wildtype littermates (**Figure 4B**). Hearts histologically were similar without any fibrosis (**Figure S5B)** and heart weight to body weight ratio were not different between genotypes (**Figure S5C)**.

**Figure 4.**
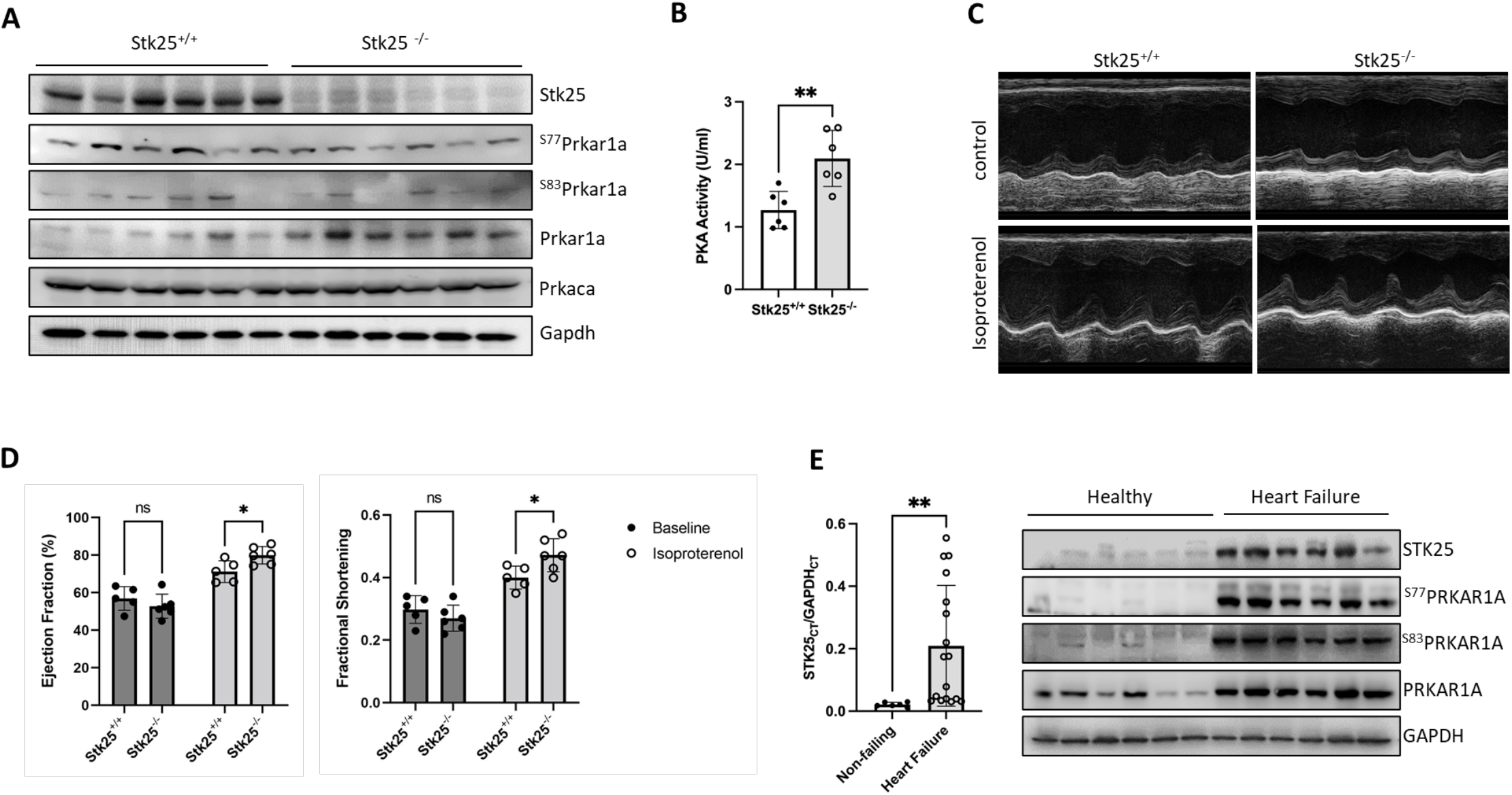
*Stk25 loss increases response to adrenergic stimulation in vivo*. A) Immmunoblot of Stk25, Prkaca, Gapdh, phospho-S77, phospho-S83 and total Prkar1a in *Stk25^+/+^* and *Stk25^-/-^* whole heart lysates. B) *Stk25^+/+^* and *Stk25^-/-^* mouse heart lysates were assessed for PKA activity *in vitro*, n=6 for each condition. C) Representative m-mode images of *Stk25^+/+^* and *Stk25^-/-^* mouse hearts stimulated with either control or isoproterenol. D) Echocardiographic measurements of ejection fraction (EF) and fractional shortening (FS) at unstimulated baseline and in response to isoproterenol, n=5 for *Stk25*^+/+^and n=6 for *Stk25^-/-^*. E) RT-PCR (left) from left ventricular myocardium of normal hearts (n=6) or heart failure (n=17) expressed as a ratio of the threshold cycle curve (Ct) of *STK25* to *GAPDH*. Immunoblot (right) of STK25 and PRKAR1A expression and phosphorylation in protein lysates from left ventricular myocardium of normal hearts or failing hearts. Bar graphs presented as mean +/- SD, *p<0.05, **p<0.01 by student t-test in 4B, repeated measures 2 way ANOVA with Sidak’s correction for multiple comparisons in 4D and Welch’s t-test in 4E.

Since Pka mediates beta adrenergic stimulation of cardiomyocyte contraction function *in* vivo, we assessed the impact of *Stk25* loss on cardiac function in response to the β1 agonist isoproterenol using echocardiography. At baseline, there were no significant differences in either contractile function or chamber dimensions between genotypes; however *Stk25^-/-^* mice displayed statistically greater individual increases in ejection fraction (EF) and fractional shortening (FS) and trends towards decreased chamber size in response to isoproterenol (**Figure 4C, 4D and S5D)**. Phosphorylation levels of downstream Pka effectors were also evaluated in response to isoproterenol and loss of *Stk25* was associated with significant increases in isoproterenol induced phosphorylation of Pln, TnI, and Ryr2 while CamKII remained unchanged (**Figure S6A and S6B**).

Because prolonged Pka activation has been associated with heart failure (Antos et al., 2001; Kushnir et al., 2010), a survival cohort of mice at 52 weeks were evaluated by echocardiography and no significant differences in either heart function or left ventricular size were found (**Figure S7**). We then investigated human heart failure samples and found that *STK25* expression was broadly *increased* in myocardial samples from patients with end stage heart failure when compared to normal heart samples (**Figure 4E and Table S1)**. Using available protein samples from this heart failure cohort, we analyzed the phosphorylation status of PRKAR1A and observed a concordant increase in STK25, PRKAR1A and phosphorylated PRKAR1A. This raises the possibility that upregulation of STK25 and PRKAR1A has a role in heart failure and may be involved in the compensatory downregulation of beta adrenergic signaling.

## Discussion

Kinases are integral components of signal transduction pathways, playing key roles in most cellular processes that are initiated by external cues. Therapeutic modulation of kinases has demonstrated success in many branches of medicine, highlighted especially by those that leverage inhibition of kinase cascades implicated in oncogenic processes. Within cardiovascular biology, kinases regulate ion handling, contractility, and metabolism, and their dysregulation is involved in many cardiovascular diseases (Clerk et al., 2007; Heineke and Molkentin, 2006; Vlahos et al., 2003). Several kinases including PKA, Ca2+ calmodulin dependent kinase, protein kinase C and PIP3 kinase have all been implicated in both physiologic and pathologic signaling (Dhalla and Muller, 2010). Despite this progress, kinase directed therapy has not been developed successfully for cardiovascular diseases, and identifying new regulators of kinase pathways is an area of unrealized therapeutic potential.

Here we present evidence that STK25 is an inhibitor of the PKA pathway, whose phosphorylation of a regulatory subunit drives inhibition of the catalytic subunit’s kinase activity. Though phosphorylation of the Type II regulatory subunits has been described in detail, little is known about phosphorylation of Type I regulatory subunits. Phosphorylation at S103 *in vitro* has been shown to be mediated by protein kinase G and interferes potentially with binding to PRKACA (Haushalter et al., 2018). Phosphorylation at S83 has been shown to modulate PRKAR1A’s association with the replication factor c complex *in vitro*, the consequence of which is unknown (Gupte et al., 2006). In the model we propose, phosphorylation at S77 and S83 increases affinity of PRKAR1A for the catalytic subunit and inhibits cAMP mediated PKA activity. This stands in contrast to the regulation of Type II subunits whose phosphorylation promotes release of the catalytic subunit and thus enhances PKA activity.

Though chronic PKA activation has been associated with progression of heart failure in animal models (Marks, 2013), knockout of *Stk25* resulted in an increase in PKA activity without evidence of heart failure. This discrepancy merits discussion. First it should be noted that heart failure mouse models overexpressing the catalytic PKA subunit generate up to 8-fold increases in PKA activity while in our model, mice experience approximately a 1.6-fold increase in PKA activity. This is in range of prior work demonstrating no heart failure phenotype with *Prkar1a* heterozygosity despite an increase in PKA activity by approximately 1.4-fold (Liu et al., 2020). Furthermore, data from human heart failure samples demonstrate *decreased* phosphorylation of PKA targets (Najafi et al., 2016) as chronic hyperstimulation of the beta-adrenergic receptor leads to desensitization and what is thought to be decreased PKA substrate activation (Ungerer et al., 1993). We observe *increased* levels of STK25 and phosphorylated PRKAR1A in heart failure samples. This would translate into suppression of PKA stimulation yet it remains to be seen whether these changes are protective (e.g. as part of a downregulation response to hyperadrenergic state of heart failure) versus maladaptive (e.g. higher levels of STK25 lead to worsening cardiac performance) or both. While, differences in either heart function or mortality are not observed with loss of *Stk25*, further longitudinal studies with overlayed chronic heart failure models are needed to determine if sustained STK25 loss is a potential therapeutic to improve cardiac function safely.

### Limitations

There are several important limitations to this study. How exactly phosphorylation of PRKAR1A alters its ability to inhibit the catalytic subunits remains unclear. We observe increased binding of S77E/S83E to the catalytic subunits in forskolin stimulated cells which implies there is either increased affinity between PRKAR1A and the catalytic subunit versus decreased affinity for cAMP by phosphorylated PRKAR1A. Another possibility is a change in PRKAR1A’s affinity for an AKAP. Though Type I regulatory subunits generally have weak affinities for AKAP proteins and display a diffuse cytoplasmic localization, the impact that phosphorylation on the assembly of a larger macromolecular regulatory complex remains to be investigated. This is particularly relevant since forskolin did not inhibit PRKAR1A and STK25 binding, yet PRKAR1A phosphorylation was diminished. This would imply that there is either inhibition of STK25 kinase activity by cAMP or a phosphatase that acts on PRKAR1A that is stimulated by cAMP. Given that we observe residual phosphorylation of S77 and S83 in both iPSC-CM and mouse knockout studies, it is likely that there are other kinases that phosphorylate these sites and regulate PKA activity. Further studies are needed to elucidate kinases and phosphatases modulate the phosphorylation status of PRKAR1A what their physiological role is in PKA activity in the heart.

In summary, loss of *STK25* leads to increased PKA activity *in vitro* and *in vivo*. We identify PRKAR1A as a new substrate for STK25 and demonstrate that phosphorylation of PRKAR1A at S77 and S83 inhibits PKA signaling. Loss of Stk25 leads to increased physiologic response to beta adrenergic stimulation of cardiac function without evidence of heart failure. Both STK25 and the phosphorylation of PRKAR1A are upregulated in heart failure and potentially represent a new pathway for downregulation of beta adrenergic signaling in the heart.

## Acknowledgements

We would like to thank the Qing Li for his surgical expertise, Erin Bush at the Columbia Genome Center for RNA-seq help, Christopher Damoci at the Columbia Mouse Imaging Core Facility and the labs of Veli Topkara, Elain Wan and Emily Tsai for reagents, technical expertise and tissue samples. B.F is supported by a grant from the NHLBI (K08HL140201), the Gerstner Foundation and the Schwartz Foundation. G.V.N is supported by grants from the NIH (UH3EB025765, P41EB027062, and R01HL076485) and NSF (NSF16478). B.W. T.N and B.L are supported by the MSTP Training Program (T32GM007367). B.L. is also supported by the NIH (F30HL145921). R.K.S is supported by 2P30 CA013696-45 Cancer Center Support Grant. M.K. is supported by a Glorney Raisbeck Fellowship Award in Cardiovascular Disease.

## Data Availability

RNA-seq data has been deposited in the Gene Expression Omnibus, NCBI: GSE195514 Proteomics data has been deposited in the PRIDE ProteomeXchange: PXD031367 Original Data can be found at: https://data.mendeley.com/datasets/t2z9bkm78h/draft?a=a7e3489f-7112-4504-94a4-33f48aa43867

## Material Availability

All unique reagents generated in this study are available from the lead contact with a completed materials transfer agreement.

## Supplementary Materials

**Figure S1.**
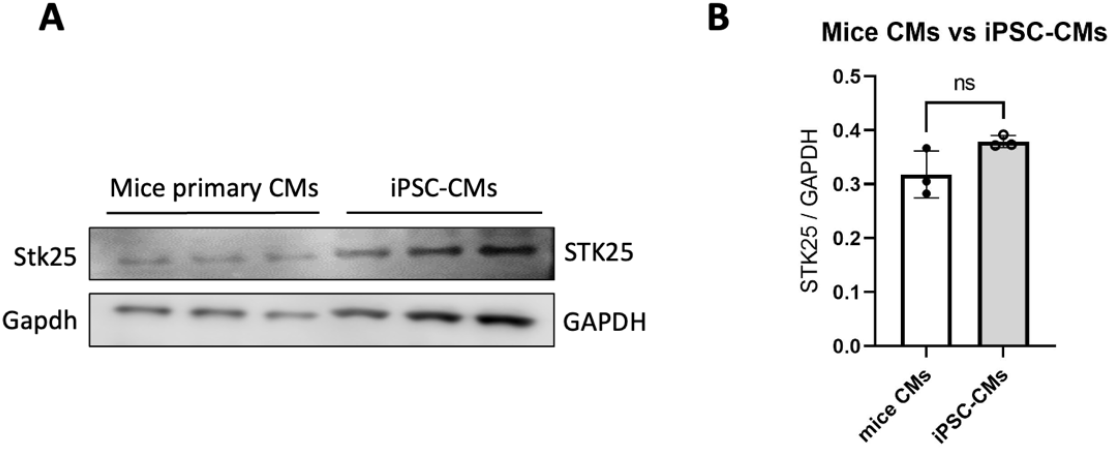
Comparison of STK25 expression in adult mice cardiomyocytes and in iPSC-CMs. A) Immunoblot and B) densitometry of STK25 and GAPDH expression in primary cardiomyocytes isolated from adult mouse heart and in iPSC-CMs. n=3 for each condition. Data presented as mean +/- SD, statistical significance was tested by student t-test in S1B.

**Figure S2.**
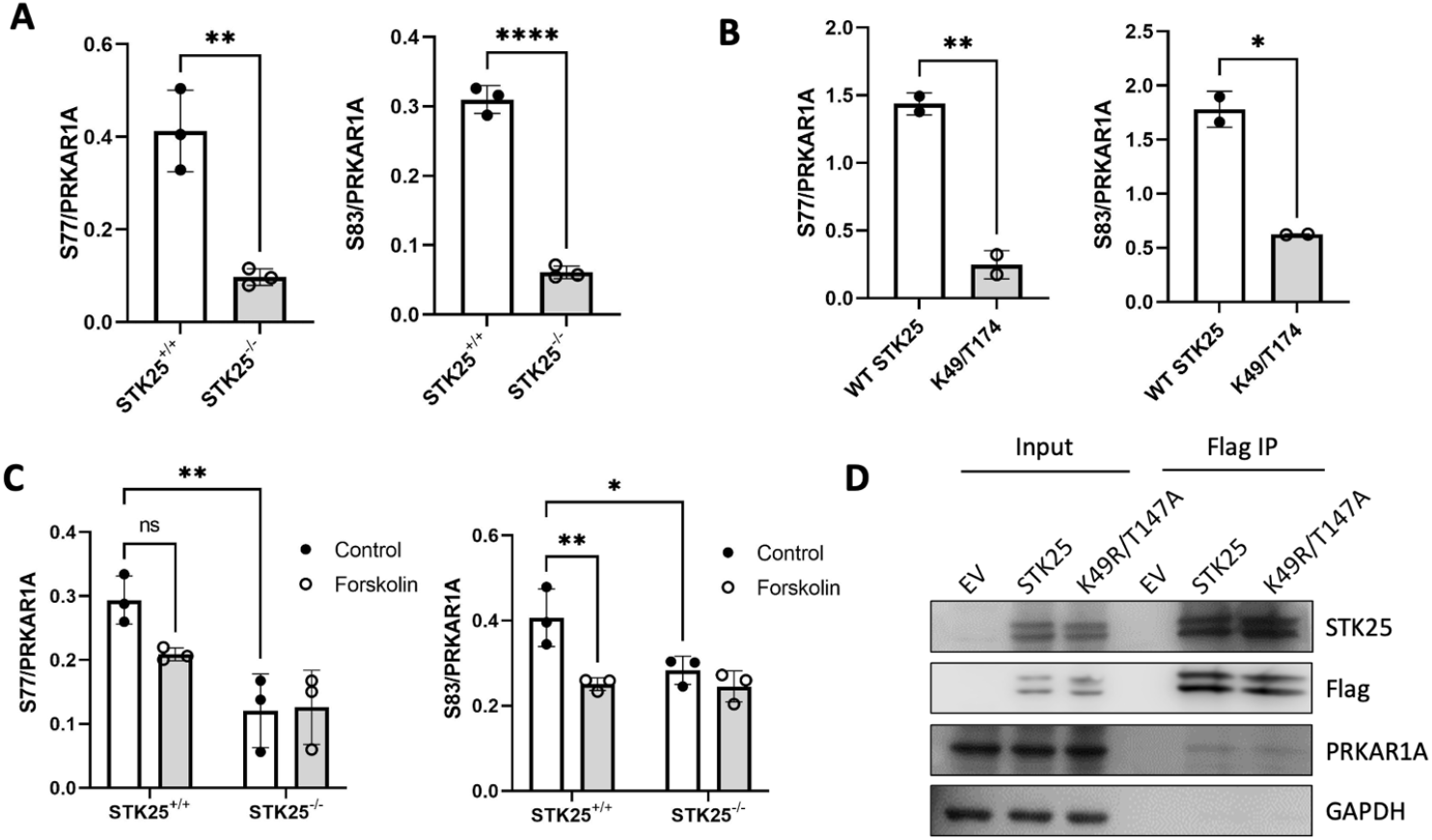
A) Densitometry analysis of phospho-S77 and S83 relative to total PRKAR1A in immunoblot shown in Figure 2A. B) Densitometry analysis of phospho-S77 and S83 relative to total PRKAR1A in immunoblot shown in Figure 2B. C) Densitometry analysis of phospho-S77 and S83 relative to total PRKAR1A in immunoblot shown in Figure 2C. D) Co-immunoprecipitation experiments of either wild type or kinase dead (K49R/T147A) mutant of STK25 in HEK293T cells with overexpressed flag tagged STK25 protein and endogenous PRKAR1A. For all bar graphs in this figure, n=3 for each condition, Data presented as mean +/- SD, *p<0.05, **p<0.01, and ****p<0.0001 by student’s t-test in S2A and S2B and two way ANOVA with Tukey’s adjustment for multiple comparisons in S2C.

**Figure S3.**
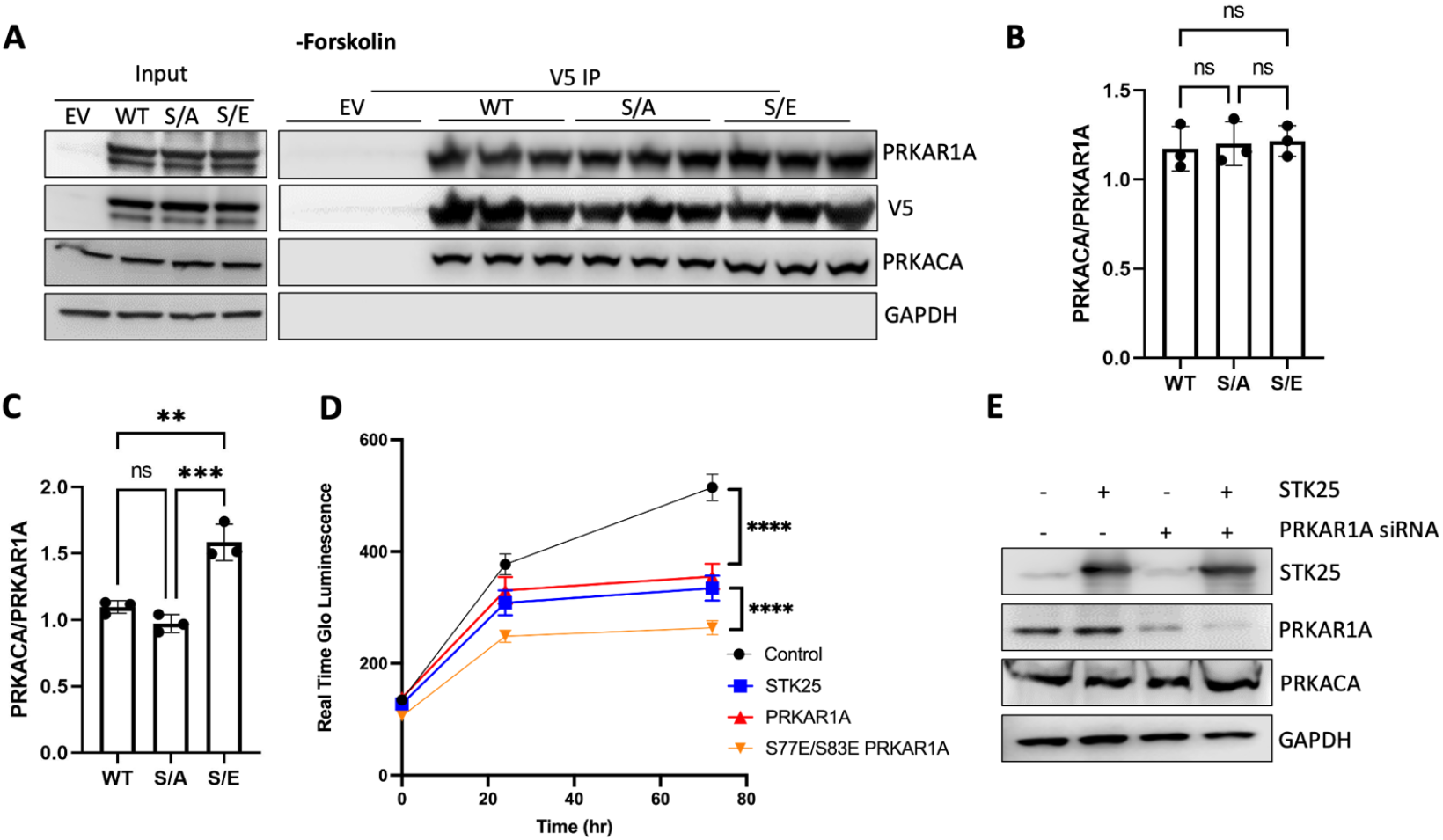
A) Co-immunoprecipitation of PRKAR1A-V5 with STK25 and PRKACA in HEK293T cells without forskolin treatment. B) Densitometry analysis of PRKACA relative to PRKAR1A in immunoblot shown in Figure S2A. C) Densitometry analysis of PRKACA relative to PRKAR1A in immunoblot shown in Figure 3A. D) HEK293T cells transfected with the indicated vectors and assessed for growth by Real Time Glo (Promega) over three days. E) Immunoblot demonstrating both overexpression of STK25 and knockdown of PRKAR1A in HEK293T cells shown in Figure 3D. For all graphs in this figure, n=3 for each condition or time point, Data presented as mean +/- SD, **p<0.01, ***p<0.001 and ****p<0.0001 one way ANOVA in S3A and S3C and two way ANOVA, all with Tukey’s adjustment for multiple comparisons.

**Figure S4:**
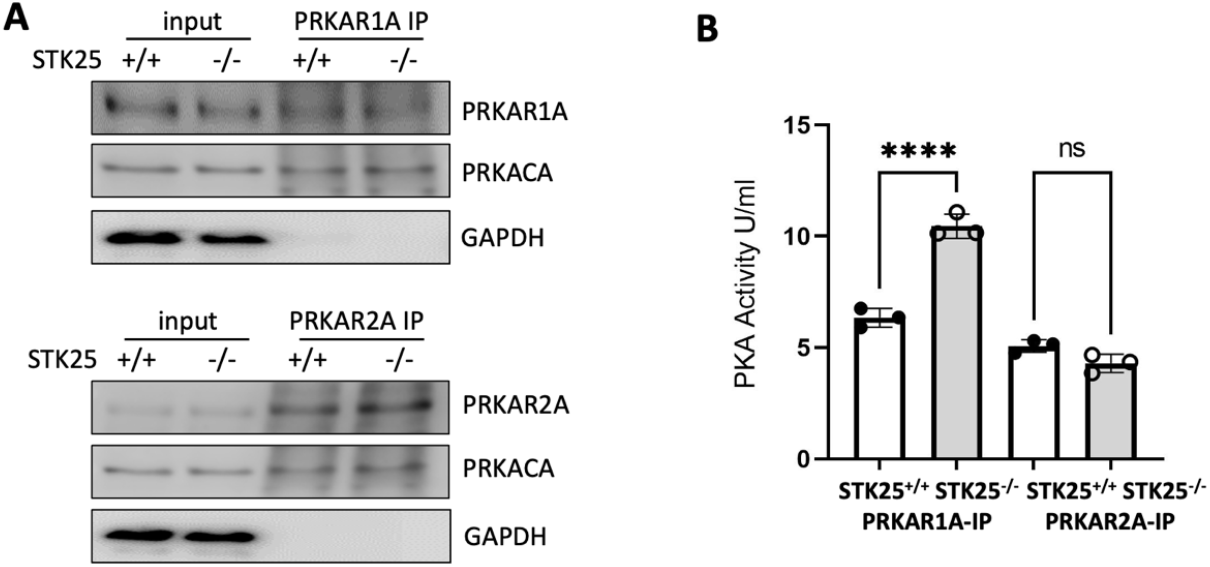
Comparison of Type I versus Type II holoenzyme activity in response to STK25 loss. A) Immunoprecipitations of PRKAR1A or PRKAR2A in STK25^+/+^ and STK25^-/-^ iPSC-CMs and immunoblotting for PRKAR1A or PRKAR2A, PRKACA and GAPDH. B) PKA activity of type I and II PKA holoenzymes precipitated from cell lysates of STK25^+/+^ and STK25^-/-^iPSC-CMs. N=3 for each condition, mean +/- SD, ****p<0.0001 two way ANOVA with Sidak’s adjustment for multiple comparisons.

**Figure S5.**
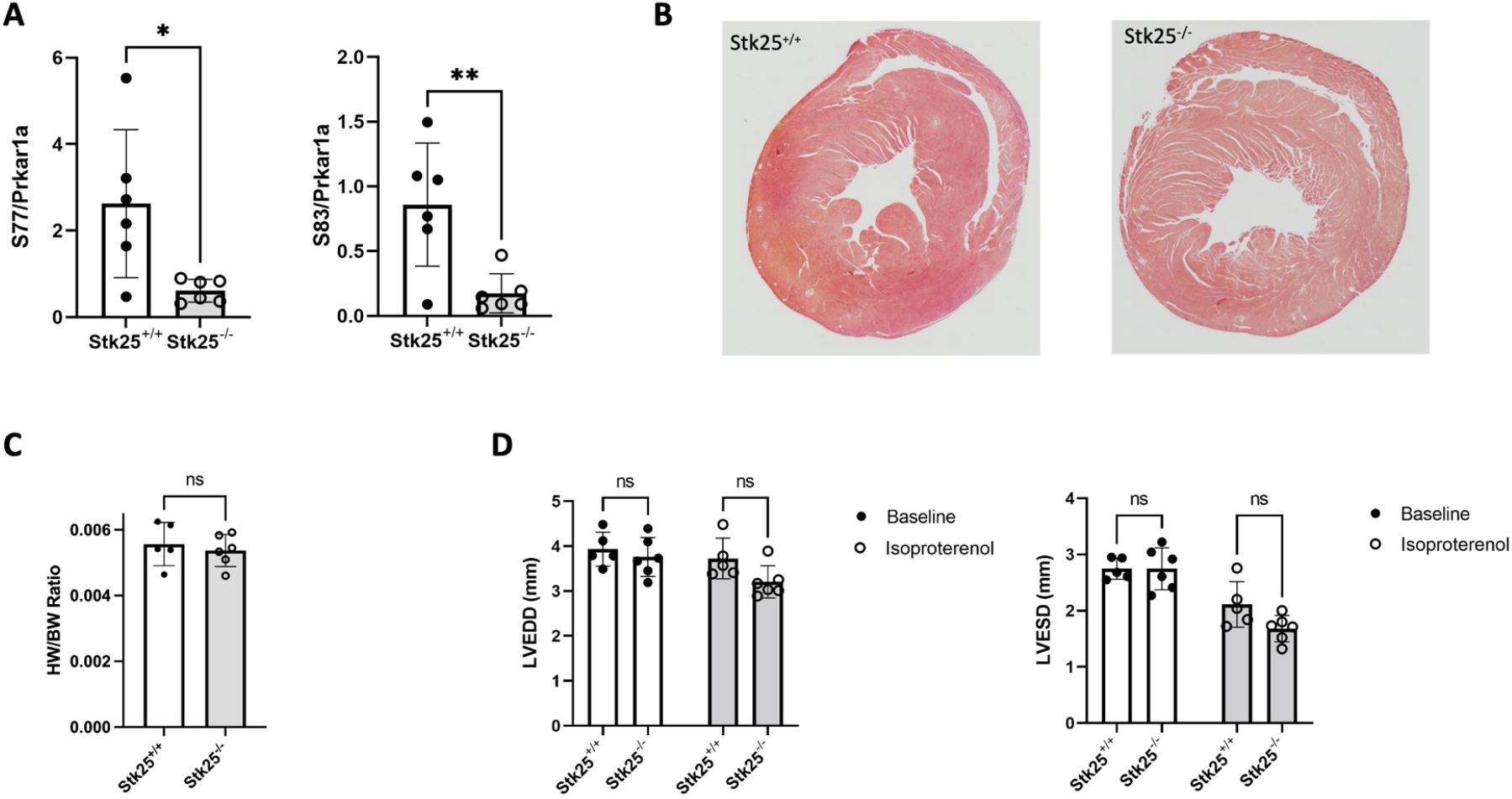
A) Densitometry analysis phospho-S77 and S83 relative to total Prkar1a in immunoblot shown in Figure 4A. B). Representative pentachrome staining of mid ventricle sections of 20-week old *Stk25^+/+^* and *Stk25^-/-^* mice. C) Heart weight (HW) to body weight (BW) ratio of mouse hearts at 20 weeks. D) Chamber measurements LVEDD (mm) and LVESD (mm) of mice at baseline and treated with isoproterenol. n=5 for *Stk25*^+/+^and n=6 for *Stk25^-/^* mice^*-*^. Data presented as mean +/- SD, statistical significance was tested with student’s t-test in S5A, Welch’s t-test in S5C, and repeated measures 2-way ANOVA with Sidak’s correction for multiple comparisons in S5D.

**Figure S6:**
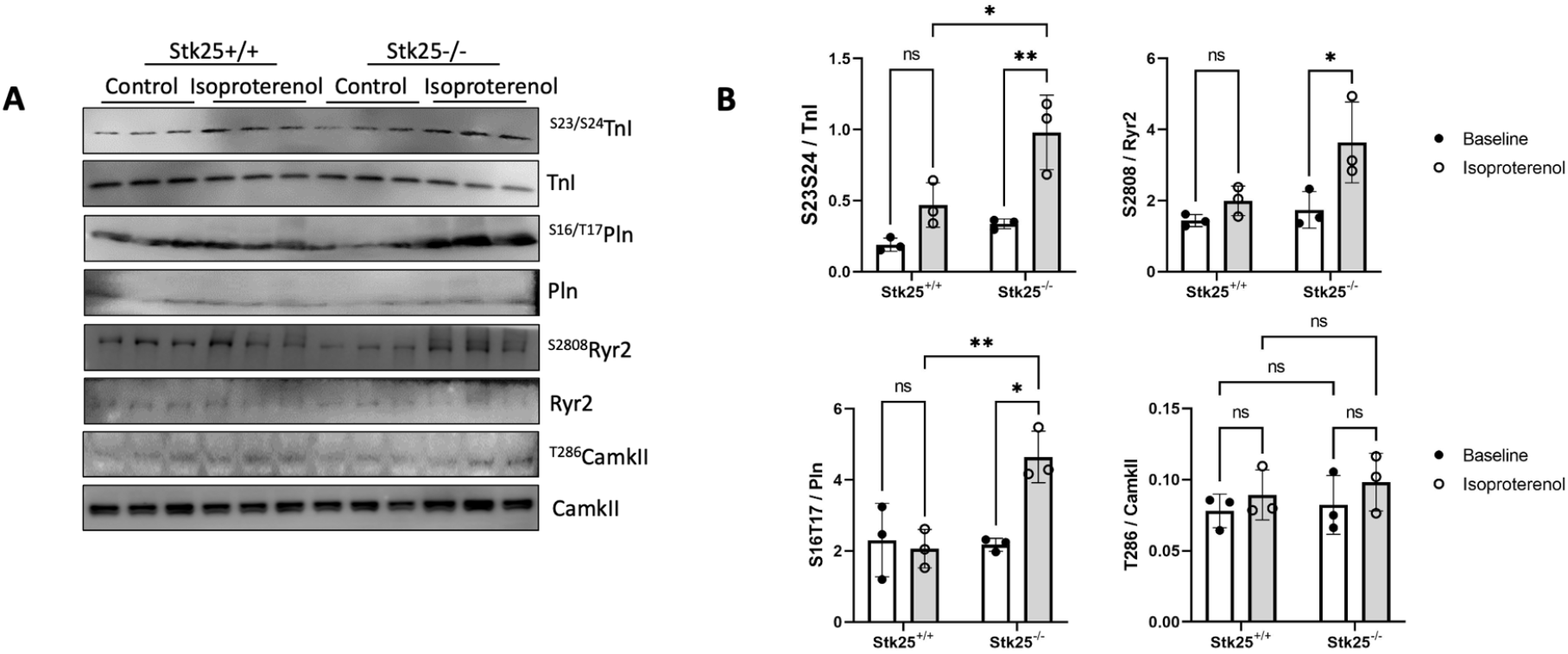
Stk25 knockout and beta-adrenergic response. A) Immunoblot of phospho-S23/S24 and total Tnl, phospho-S16/T17 and total Pln, phospho-S2808 and total Ryr2, phospho-T286 and total CamkII in *Stk25^+/+^* and *Stk25^-/-^* whole heart lysates with and without isoproterenol injection. B) Densitometry analysis of phospho-S23/S24 relative to total TnI, phospho-S16/T17 relative to total Pln, phospho-S2808 relative to total Ryr2, phospho-T286 relative to total CamkII in immunoblot shown in Figure S6C, n=3 for each condition. Data presented as mean +/- SD, *p<0.05, **p<0.01, statistical significance was tested with two way ANOVA with Tukey’s correction for multiple comparisons.

**Figure S7:**
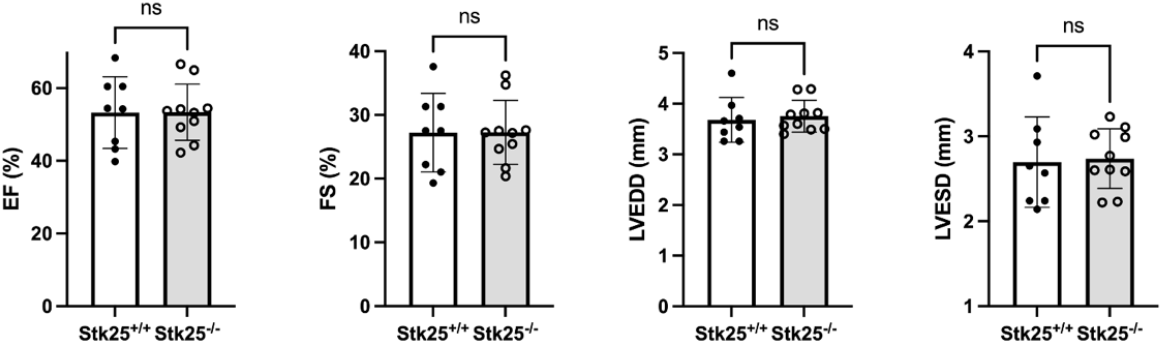
Cardiac function at one year in Stk25-/-mice. Echocardiographic measurements of LV function and size in a survival cohort at 52 weeks for Stk25+/+ (n=8) and Stk25-/-(n=10). Data presented as mean +/- SD, statistical significance was tested with a Welch’s t-test.

**Table S1:** Patient characteristics of end stage heart failure samples.

| Patient Characteristics (n=17) |  |  |
| --- | --- | --- |
|  | Age (SD) | 49.47 (12.8) |
|  | Gender (Male%) | 82.4 |
|  | BMI (SD) | 24.4 (3.3) |
| Cardiomyopathy |  |  |
|  | Non-ischemic (%) | 70.6 |
|  | Ischemic (%) | 29.4 |
| Comorbidities |  |  |
|  | Hypertension (%) | 17.6 |
|  | Hyperlipidemia (%) | 29.4 |
|  | Diabetes(%) | 23.5 |
| Heart Function |  |  |
|  | Ejection Fraction (%) | 17.9 |
|  | LVAD | 88.23 |
|  | Inotrope | 35.3 |
| Medical Therapy |  |  |
|  | ACEi/ARB/ARNI | 29.4 |
|  | Beta Blocker | 76.5 |
|  | MRA | 52.9 |
|  | Anticoagulation | 88.23 |
|  | Antiarrhythmic | 41.2 |

Abbreviations: SD, standard deviation; BMI, body mass index; LVAD, left ventricular assist device; ACEi, angiotensin converting enzyme inhibitor; ARB, angiotensin II receptor blocker; ARNI angiotensin II receptor blocker and neprilysin inhibitor; MRA, mineralocorticoid receptor antagonist.

